# A Multimodal Vision Transformer for Interpretable Fusion of Functional and Structural Neuroimaging Data

**DOI:** 10.1101/2023.07.14.549002

**Authors:** Yuda Bi, Anees Abrol, Zening Fu, Vince D. Calhoun

## Abstract

Deep learning models, despite their potential for increasing our understanding of intricate neuroimaging data, can be hampered by challenges related to interpretability. Multimodal neuroimaging appears to be a promising approach that allows us to extract supplementary information from various imaging modalities. It’s noteworthy that functional brain changes are often more pronounced in schizophrenia, albeit potentially less reproducible, while structural MRI effects are more replicable but usually manifest smaller effects. Instead of conducting isolated analyses for each modality, the joint analysis of these data can bolster the effects and further refine our neurobiological understanding of schizophrenia. This paper introduces a novel deep learning model, the multimodal vision transformer (MultiViT), specifically engineered to enhance the accuracy of classifying schizophrenia by using structural MRI (sMRI) and functional MRI (fMRI) data independently and simultaneously leveraging the combined information from both modalities. This study uses functional network connectivity data derived from a fully automated independent component analysis method as the fMRI features and segmented gray matter volume (GMV) as the sMRI features. These offer sensitive, high-dimensional features for learning from structural and functional MRI data. The resulting MultiViT model is lightweight and robust, outperforming unimodal analyses. Our approach has been applied to data collected from control subjects and patients with schizophrenia, with the MultiViT model achieving an AUC of 0.833, which is significantly higher than the average 0.766 AUC for unimodal baselines and 0.78 AUC for multimodal baselines. Advanced algorithmic approaches for predicting and characterizing these disorders have consistently evolved, though subject and diagnostic heterogeneity pose significant challenges. Given that each modality provides only a partial representation of the brain, we can gather more comprehensive information by harnessing both modalities than by relying on either one independently. Furthermore, we conducted a saliency analysis to gain insights into the co-alterations in structural gray matter and functional network connectivity disrupted in schizophrenia. While it’s clear that the MultiViT model demonstrates differences compared to previous multimodal methods, the specifics of how it compares to methods such as MCCA and JICA are still under investigation, and more research is needed in this area. The findings underscore the potential of interpretable multimodal data fusion models like the MultiViT, highlighting their robustness and potential in the classification and understanding of schizophrenia.

## I. Introduction

Deep learning has emerged as a rapidly evolving field in recent years, significantly contributing to addressing various challenges in computer vision and image processing tasks. The success of Convolutional Neural Networks (CNNs) and their numerous adaptations have resulted in the widespread adoption of deep learning techniques in neuroimaging research [1]. Numerous studies have employed 3D CNN models to predict intricate brain disorders such as schizophrenia, with the majority of these studies concentrating on unimodal data [2], [3], [4]. In computer vision, a novel model called the Vision Transformer (ViT) [5] has recently gained prominence. ViT is an extension of the transformer model, initially developed for natural language processing [6]. ViTs hold the potential to surpass traditional CNN models in various computer vision tasks, including image classification [7], image segmentation [8], and object detection [9]. ViT can effectively substitute CNN-based models due to its superior performance on large-scale image datasets; the self-attention mechanism in ViTs results in larger receptive fields and better encoding of spatial relationships among different features and preserves long-range dependency, offering a structural advantage over conventional CNN models. Nevertheless, due to privacy concerns and costly data collection processes, the scarcity of data in medical imaging has prompted some researchers to explore hybrid CNN-Transformer models to tackle medical imaging challenges [10], [11]. ViT’s performance is contingent upon the size of the dataset, and hybrid models may help surmount this limitation.

The objective of this study is to devise an interpretable and scalable deep learning model capable of integrating structural magnetic resonance imaging (sMRI) and functional magnetic resonance imaging (fMRI) data, based on the assumption that multimodal MRI data have mutual information that can enhance disease prediction. To capitalize on the information from both modalities, parallel data input channels are established within the proposed pipeline. Several renowned deep learning models are assessed as training bases, encompassing 3D convolutional neural networks (3DCNNs), vision transformers (ViTs), and traditional machine learning methods. Building on multimodal datasets, a multimodal ViT framework, dubbed **MultiViT**, is proposed as a benchmark model for predicting and diagnosing schizophrenia with increased accuracy. The framework comprises two parallel multiscale vision transformer encoders connected by cross-attention encoder layers, facilitating the mutual sharing of information across two or more modalities. The pipeline can generate anatomical brain saliency maps from feature extractions following the cross-attention layer, accentuating regions that exhibit distinctive patterns for schizophrenia and healthy subjects. Functional network connectivity (FNC) data can also be employed to obtain highlighted matrix regions associated with a specific prediction, revealing functional patterns corresponding to structural changes.

The proposed MultiViT model is applied to study schizophrenia, a severe psychiatric disorder characterized by intricate alterations in brain anatomy and function and altered connections between cortical regions. Neuroimaging data can yield valuable insights into mental illnesses such as schizophrenia by exposing brain regions and connections correlated with the disorder [12], [13]. Prior studies have utilized various neuroimaging modalities and cutting-edge artificial intelligence models to investigate schizophrenia. The most consistent findings using sMRI data disclose reduced gray matter volumes in brain regions essential for processing auditory information, short-term memory, and decision-making [14]. Moreover, functional fMRI observations have effectively identified schizophrenia based on blood oxygen level dependent (BOLD) signals. Although some studies have explored the relationships between structural changes and functional abnormalities in schizophrenia [15], [16], limited research concentrates on simultaneously capturing and visualizing intricate relationships between functional network connectivity and gray matter data.

In the present study, several noteworthy contributions have been made, as enumerated below:

1. A novel multimodal ViT framework has been devised, which effectively amalgamates structural and functional neuroimaging data. This is achieved by employing cross-attention techniques to retain the mutual information between the two modalities.
2. Rigorous validation of the proposed pipeline has been conducted against various benchmarks, encompassing unimodal and multimodal baselines. These results substantiate the efficacy of the newly developed pipeline design.
3. By implementing state-of-the-art attention-driven saliency mapping on structural and functional neuroimaging data, the research identifies brain regions that strongly correlate with schizophrenia. Consequently, this study bridges the gap between structural brain regions and their potential functional implications.

## II. Related Works

Models trained to utilize a singular data type have been observed to enhance multimodal models through asymmetric data fusion [17]. Neuroimaging techniques, including magnetic resonance imaging (MRI), electroencephalogram (EEG), positron emission tomography (PET), diffusion tensor imaging (DTI), and others, play an essential role in the accurate diagnosis of brain diseases [17]. In deep learning, multimodal research endeavors to integrate computer vision with auditory and textual data to refine the precision of computer vision tasks, such as image classification and object detection [18]. Likewise, researchers strive to amalgamate different medical imaging modalities to improve disease prediction outcomes. For example, Parvathy *et al*. [19] propose an algorithm capable of extracting meaningful features from diverse modalities and employing feature decomposition techniques to mitigate complexity. Lei *et al*. [20] extract characteristics from structural and functional MRI data and classify schizophrenia using classical machine learning techniques, such as support vector machines (SVM). More recently, some studies are using SOTA deep learning models, such as Sarraf *et al*. [21] uses optimized vision transformers to predict various stages of Alzheimer’s disease.

Multimodal vision transformers (ViTs) have been employed in a variety of computer vision and natural language processing tasks [22] for multi-modality analysis. Shvetsova *et al*. [23], for instance, developed a multi-modal fusion ViT model capable of exchanging information between multiple modalities, including video, audio, and text. Wei *et al*. [24] proposed a cross-attention network that can exploit intra-modality relationships within each modality, ensuring robustness for image and sentence matching. However, the majority of ViT models utilized in neuroimaging research are unimodal rather than multimodal. Singla *et al*. [25] also developed a 3D-ViT model for gender prediction tasks using structural MRI data, achieving over 0.9 Area Under the Curve (AUC) on the Adolescent Brain Cognitive Development (ABCD) dataset. Nevertheless, recent studies [26][27] have demonstrated that a 3D convolutional neural network (CNN) architecture can achieve a 0.95 AUC on the ABCD dataset for gender prediction, suggesting that a pure ViT model may not surpass 3D CNN in specific tasks when handling complex neuroimaging data. Dai *et al*. [10] employed a CNN-Transformer hybrid model to classify brain images from multiple datasets, exhibiting strong performance on Alzheimer’s disease-related datasets. Venugopalan *et al*. [28] proposed a framework for multimodal data fusion of MRI imaging data, electronic health records (EHRs), and single nucleotide polymorphism (SNP) data to enhance Alzheimer’s disease prediction, ultimately concluding that their deep learning framework outperforms single-modality models. Oh *et al*. [29] proposed a 3D CNN model for schizophrenia disease prediction based on structural MRI data, achieving an AUC score of approximately 0.71. Functional MRI data were utilized by [30] to create a modified 3D Visual Geometry Group (VGG) model for schizophrenia disease classification, attaining an accuracy of 84.3%. However, these studies present several limitations, such as diminished detection rates, limited model interpretability, and a lack of focus on global dependencies in the data.

## III. Methods

Our methodology comprises the following stages: Initially, we conduct preprocessing on the input data (structural MRI and functional MRI) utilizing established pipelines. This involves generating 3D gray matter images from structural MRI using the unified segmentation model in the SPM toolbox and computing functional MRI features employing the NeuroMark pipeline[31], a fully automated spatially constrained independent component analysis. Subsequently, we construct a 2D static FNC matrix. Secondly, we concurrently analyze the structural and functional MRI modalities employing a multi-modal deep learning framework that encodes the data through specialized architectures. We assessed various multimodal pipelines, including 3DCNN-3DCNN, 3DViT-3DViT, 3DViT-2DCNN, and 3DViT-MLP. Upon training the deep learning baselines, we established the MultiViT model as a benchmark that outperforms all other deep learning and machine learning variations. Finally, we developed an interpretable architecture capable of generating attention maps from 3D structural MRI and 2D FNC data. These maps can be utilized to identify potential connections between the structural and functional MRI data concerning schizophrenia. Figure 1 delineates the general outline of our research.

**Fig. 1.**
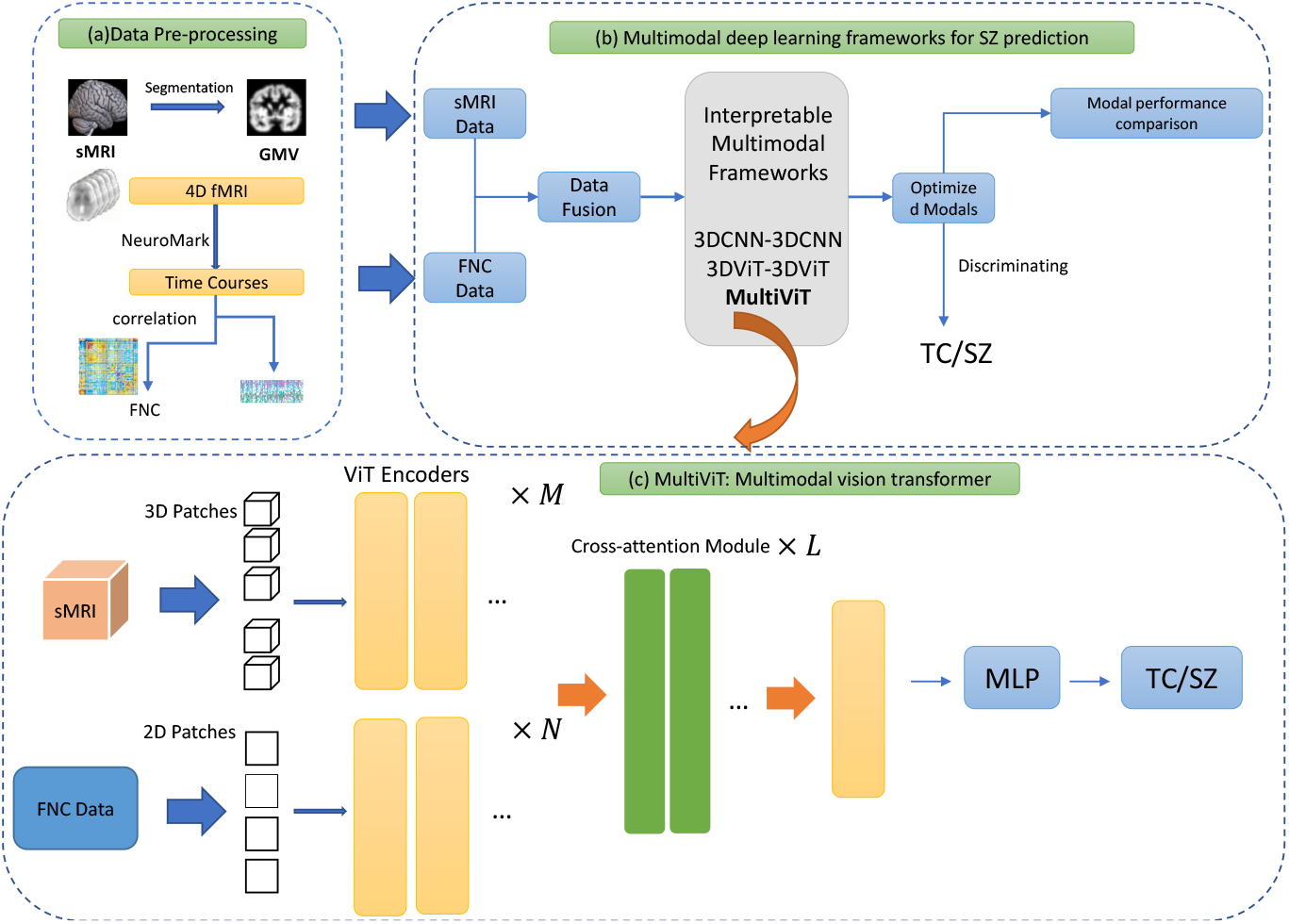
The general pipeline:(a) Data pre-processing module for sMRI segmentation and fMRI pre-processing using NeuroMark pipeline[31] to FNC and time courses; (b) General training process for multimodal deep learning frameworks; (c) The proposed MultiViT architecture.

### A. Data and Pre-processing

In this study, we employed two datasets related to clinical research on schizophrenia, Table I shows the subjects’ information of datasets. The first dataset was compiled from three distinct studies, namely fBIRN (Functional Imaging Biomedical Informatics Research Network) with seven sites, MPRC (Maryland Psychiatric Research Center) with three sites [there are some duplicate subjects between sites, do you remove them?], and COBRE (Center for Biomedical Research Excellence) with one site. This resulted in a total of 827 participants, including 477 control individuals (mean age: 38.76 ± 13.39, 213 females, 264 males) and 350 schizophrenia individuals (mean age: 38.70 ± 13.14, 96 females, 254 males). The fBIRN data was collected using the same parameters for resting-state fMRI (rsfMRI) at all sites, employing a standard gradient echo-planar imaging (EPI) sequence with a repetition time (TR) and echo time (TE) of 2000/30 ms, a voxel spacing size of 3.4375 × 3.4375 × 4 mm, and a field of view (FOV) of 220 × 220 mm. The data was collected using 6 Siemens Tim Trio 3-Tesla scanners and 1 General Electric Discovery MR750 3.0 Tesla scanner. For the COBRE data, rsfMRI images were collected using a standard EPI sequence with a TR/TE of 2000/29 ms and a voxel spacing size of 3.75 × 3.75 × 4.5 mm a FOV of 240 × 240 mm, using a 3-Tesla Siemens Tim Trio scanner. The MPRC data was collected using three different 3-Tesla Siemens scanners, including the Siemens Allegra, Trio, and Tim Trio [32].

**TABLE I.** Dataset details.

| Dataset | Age | Gender(female/male) | Race |
| --- | --- | --- | --- |
| Dataset1 | TC: $38.76 \pm 13.39$<br>SZ: $38.70 \pm 13.14$ | TC: 213/264 (44%)<br>SZ: 96/254 (27%) | No record(from US hospitals) |
| Dataset2 | TC: $29.81 \pm 8.68$<br>SZ: $28.98 \pm 7.63$ | TC: 167/159 (51%)<br>SZ: 229/260 (45%) | Chinese ethnic Han<br>(from Chinese hospitals) |

The second dataset consisted of 815 participants, collected from seven Chinese hospitals, including Peking University Sixth Hospital, Beijing Huilongguan Hospital, Xinxiang Hospital Simens, Xinxiang Hospital GE, Xijing Hospital, Renmin Hospital of Wuhan University, and Zhumadian Psychiatric [33] Hospital. The participants included 326 control individuals (mean age: 29.81 ± 8.68, 167 females, 159 males) and 489 schizophrenia individuals (mean age: 28.98 ± 7.63, 229 females, 260 males), all of whom were Han Chinese. The resting-state fMRI data were collected using three different 3-Tesla scanners across the seven sites, including the Siemens Tim Trio, Siemens Verio, and Signa HDx GE Scanner. The participants were instructed to lie still and relax while remaining awake and calm.

The preprocessing of fMRI included slice timing correction, realignment, normalization to the EPI template, and finally smoothing with a 6 mm kernel. Details of preprocessing steps can be found in our previous studies [31]. Moreover, the sFNC data was calculated using cross-correlation among fMRI time series obtained through independent component analysis (ICA), employing a fully automated spatially constrained ICA algorithm and the neuromark_fMRI_1.0 template as spatial priors. The sMRI data were preprocessed using a voxel-based morphometry pipeline and modulated by the Jacobian of the spatial transform to produce voxelwise gray matter volume data.

### B. Transformer

The transformer model, introduced by Vaswani et al. (2017) [6], has had a profound impact on the field of natural language processing and, more broadly, deep learning. This innovative architecture overcomes the limitations of traditional RNNs and CNNs in capturing long-range dependencies and parallelization. Central to the transformer model is the concept of self-attention mechanisms, which enable the model to weigh the importance of each token in a sequence while considering the relationships between tokens at various positions. This novel approach has led to significant advancements in tasks such as machine translation, text summarization, and question-answering. Moreover, the transformer model has served as the foundation for the development of numerous state-of-the-art pre-trained models, such as BERT, GPT, and T5, which have established new performance benchmarks across a wide range of NLP tasks. Consequently, the transformer model has become an indispensable building block in the realm of deep learning, inspiring new research directions and applications beyond natural language processing.

The vision transformer (ViT) represents a notable extension of the transformer model from natural language processing to the domain of computer vision. Introduced by Dosovitskiy et al. (2020) [5], ViT adapts the self-attention mechanisms of the transformer to process image data by dividing input images into non-overlapping patches and linearly embedding them into a sequence of tokens. As a result, ViT is capable of capturing intricate spatial relationships and long-range dependencies within images. This groundbreaking approach has demonstrated exceptional performance on various computer vision tasks, including image classification, object detection, and segmentation, frequently outperforming traditional CNNs. The success of ViT is largely attributable to its ability to utilize large-scale image datasets for pre-training, which allows the model to learn more expressive and transferable visual representations. Consequently, the vision transformer has emerged as a potent and versatile instrument in the field of computer vision, spurring novel research avenues and the development of hybrid models that combine the strengths of transformers and CNNs to address a wide range of visual tasks.

Figure 2(a) illustrates the fundamental component of ViTs: the ViT encoder, which employs self-attention to encode patch embeddings from the original input images. ViTs share certain similarities with traditional CNNs, with the exception that they utilize multiple ViT encoders instead of a sequence of convolutional layers. Analogously, the multi-head self-attention mechanism is comparable to multiple kernels in a convolutional layer, effectively capturing and encoding the essential features of the input images.

**Fig. 2.**
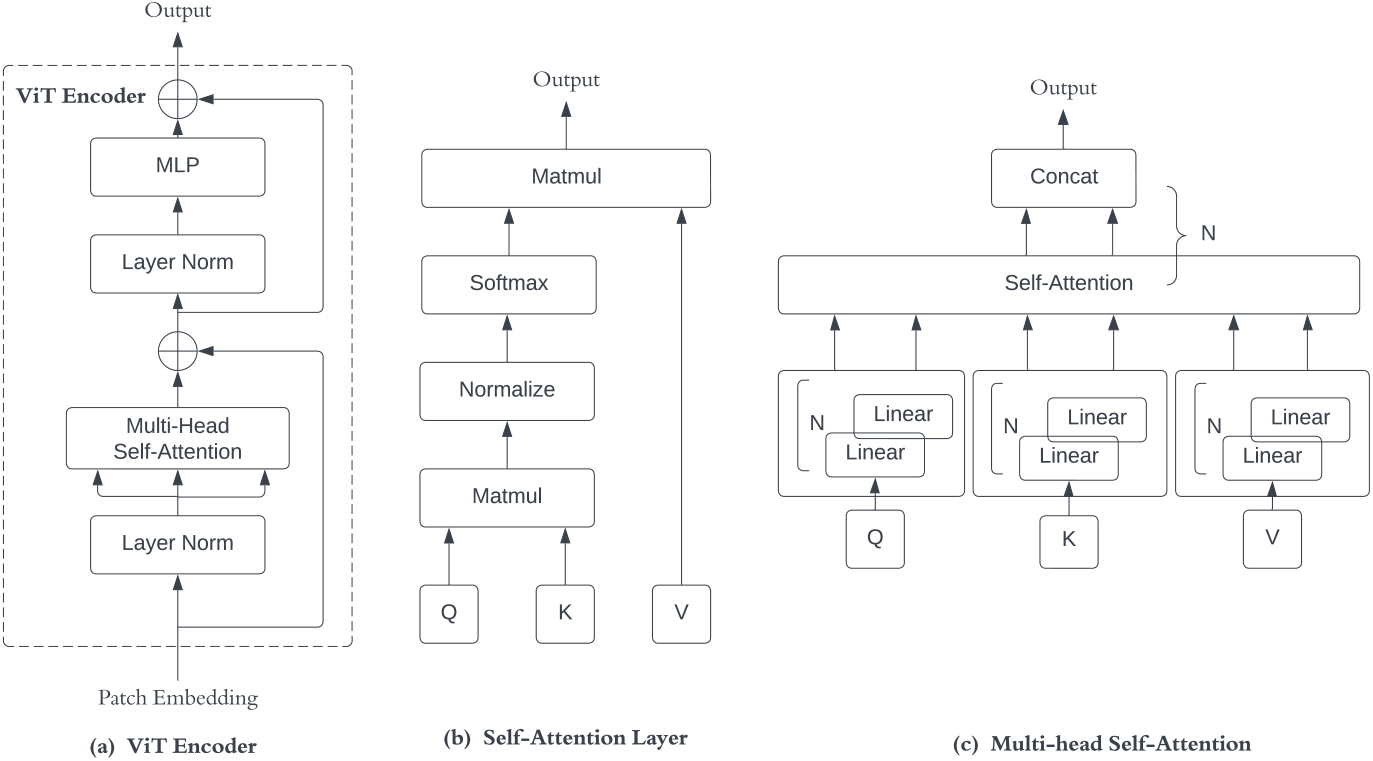
(a) The basic architecture of vision transformer encoder (ViT Encoder), (b) Self-Attention mechanism, and (c) Multi-Head Self-Attention Layers inside of ViT Encoder.

### C. Self-Attention

The pioneering application of the self-attention mechanism (SA) [34] was in the natural language processing (NLP) domain, where it was utilized to determine the level of emphasis of each word in an input sequence. Subsequently, the ViT models have expanded the use of SA to image embeddings. Given an input image embedding 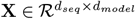, three trainable weight matrices 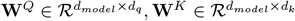, and 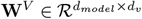 are utilized to project **X** into query matrix 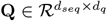, key matrix 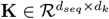, and value matrix 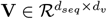.

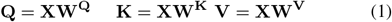

Then, we have the self-attention(SA) function that calculates the score matrix:

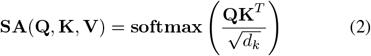

Where *d*_*q*_ = *d*_*k*_, which is the dimension of the query and key matrix. Figure 2(b) shows the self-attention steps. In the ViT models, multi-head self-attention separates the original input embeddings into multiple heads, then computes the concatenation from each head’s self-attention result. This can be represented as:

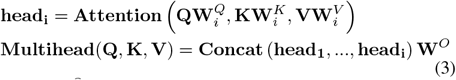

Where 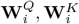, and 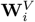 are trainable weight matrices that project the original query, key, and value, respectively, into linearly projected versions in each head *head*_*i*_. The trainable weight matrix **W**^*O*^ performs a linear projection to the final result after the concatenation of each *head*_*i*_.

### D. Cross-Attention

Cross-attention is a pivotal mechanism in transformer-based deep learning architectures that facilitates the effective integration of information from multiple input modalities. Introduced by Vaswani et al. (2017) [6], cross-attention operates by enabling a transformer model to attend to and weigh the importance of tokens or features from distinct input sources while simultaneously considering their relationships. This mechanism allows the model to effectively fuse and process heterogeneous data, such as text, audio, and images [35], or structural and functional brain imaging data, by leveraging the self-attention mechanisms inherent in transformer architectures. Cross-attention has been instrumental in advancing the performance of multimodal deep learning models, as it encourages the efficient exchange of information across different modalities [36], leading to a more comprehensive understanding of the data and improved predictive capabilities. Consequently, cross-attention has emerged as a vital component in the development of state-of-the-art models for diverse applications spanning natural language processing, computer vision, and biomedical research, among others [37].

Given two image embeddings **X** and **Y**, similar to self-attention, we can query **Q**_*X*_ and key **K**_*X*_ from embedding **X**, but **V**_*Y*_ from embedding **Y**, so that:

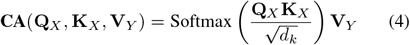

### E. MultiViT

In this section, we introduce an innovative ViT model capable of processing both structural and functional MRI data using a cross-attention mechanism. The input structural MRI image is denoted as **x** ∈ ℛ^*B×C×L×W×H*^, where *B* represents the batch size, *C* corresponds to the number of channels, and *L, W*, and *H* signify the length, width, and height of **x**, respectively. The input is partitioned into *n* 3D patches with dimensions 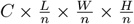 and subsequently processed through *M* ViT Encoder layers to generate an image embedding **X** ∈ ℝ^*seq*1*×model*^.

Analogously, the input functional MRI data, represented as 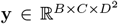, is divided into *m* 2D square patches of dimensions 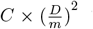. These patches undergo a concurrent embedding process via *N* ViT Encoder layers, yielding an image embedding **Y** ∈ ℝ^*Seq*2*×model*^. The cross-attention module, which comprises *L* cross-attention layers, facilitates efficient collaboration and information exchange between **X** and **Y**. Ultimately, the two components are concatenated and passed through a multilayer perceptron module (MLP) to estimate a prediction score.

### F. Attention Map

Our proposed model exhibits the capacity to generate high-level attention-based saliency maps for both structural and functional MRI data, thereby promoting the identification of relationships between structural and functional patterns in schizophrenia. Traditional CNN-based models derive saliency maps from gradients present in the activation layers of the model, emphasizing potential regions of interest within the original structural brain images. In contrast, ViT-based models produce attention maps by leveraging saliency maps from their self-attention and cross-attention layers.

Vision transformer-based attention maps present several advantages over conventional gradient-based saliency maps derived from CNN models. ViTs employ self-attention mechanisms to process entire images in parallel, taking into account the relationships among all image components. This enables the inclusion of supplementary contextual information in the prediction process, which could potentially result in improved accuracy. Additionally, ViTs are capable of capturing long-range dependencies between various regions of an image, whereas CNNs generally focus on local features and may neglect crucial long-range information. Lastly, attention maps from ViTs are more interpretable than their CNN-derived saliency map counterparts, as they underscore the areas of an image on which the model concentrates during prediction. This can yield valuable insights into the structural attributes of brain function and potentially relevant regions associated with schizophrenia.

Moreover, our multimodal framework, MultiViT, can concurrently generate attention maps from structural MRI data and FNC matrices upon the activation of cross-attention layers, thereby enabling the model to discern interpretable connections between structural and functional neuroimaging data. This may facilitate a more comprehensive understanding of schizophrenia. The cross-attention layers serve a critical function in data fusion, establishing a communication conduit between the two modalities. As evidenced by some studies, this can culminate in significant enhancements in accuracy.

## IV. Experiment

### A. Experimental Setup

#### Feature extraction

In our research, we utilized data from multiple multi-site studies on schizophrenia, including individuals scanned with a 3T MRI scanner (N=2130). The sMRI data is three-dimensional with dimensions of (121, 145, 121), and the sFNC data, calculated from 53 regions of interest from fMRI, is two-dimensional with dimensions of (53, 53). Using Torchio, we resized the sMRI and sFNC data to (120, 140, 120) and (54, 54) to fit our model due to the ViT’s input characteristics. This open-source Python library efficiently loads, preprocesses, and augments 3D medical imaging data. Additionally, we augmented our data using various strategies such as RandomCrop, RandomAffine, RandomFlip, and Add GuassianNoise, provided by Torchio. Our experiments used the PyTorch deep learning framework and NVIDIA RTX v100 GPUs.

#### Models design

Models design. To demonstrate the novelty and efficacy of the MultiViT model, we have designed a three-tiered experimental approach based on various baselines. Initially, we opted to employ a range of unimodal models, encompassing traditional machine learning techniques (e.g., support vector machines) and deep learning approaches (CNNs and ViTs) for structural and functional MRI data. These unimodal models form our first-level baseline. Subsequently, we explored multimodal models, such as 3D CNN to CNN (3DCNN-CNN) concatenation and 3D ViT to ViT (3DViT-ViT) concatenation. Finally, we proposed the MultiViT model, composed of a 3D ViT and a 2D ViT connected via a cross-attention mechanism. In contrast to simple concatenation, the cross-attention connection employed by the MultiViT model offers improved mutual information advantages.

#### Training and Evaluation

All pipelin baseline models and MultiViT, were trained to techniques such as AdamW optimization, StepLR scheduler, and a 30-epoch warmup. An 8:1:1 split was employed to select the training, validation, and testing sets. A 10-fold cross-validation approach was employed for each model to ensure accurate results. The initial hyperparameters for MultiViT training were set to a learning rate of 3 ×10^4^ and a weight decay of 1 ×10^3^, totaling 200 epochs. The models were evaluated using general accuracy, balanced accuracy, AUC, F1 score, and precision. The number of parameters for each model is also reported to summarize their complexity.

### B. Results

#### Model Performance

In this part, we summarize model performance based on validation and testing metrics for baselines and MultiViT. Table 2 shows our models’ performance, including unimodal baselines, multimodal baselines, and MultiViT. It shows that MultiViT significantly improves over other baseline models based on balanced accuracy, AUC, and F1 score. Figure 3 shows that MultiViT has superior learning performance due to its potential quick model convergence and stability over two baselines.

**TABLE II.**
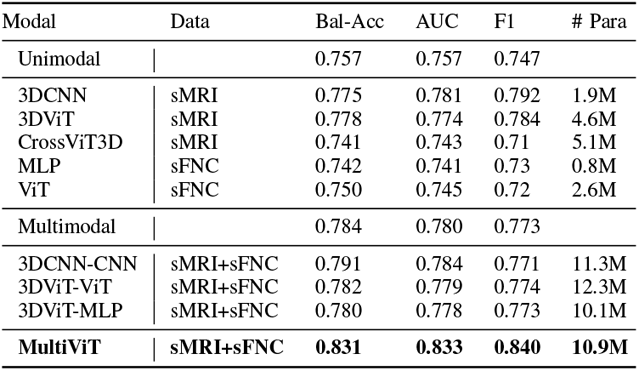
Testing results of each baseline and MultiViT: we calculate average testing metrics for both unimodal and multimodal baselines, then compare them with our new multimodal pipeline (MultiViT).

| Modal | Data | Bal-Acc | AUC | F1 | # Para |
| --- | --- | --- | --- | --- | --- |
| Unimodal |  | 0.757 | 0.757 | 0.747 |  |
| 3DCNN | sMRI | 0.775 | 0.781 | 0.792 | 1.9M |
| 3DViT | sMRI | 0.778 | 0.774 | 0.784 | 4.6M |
| CrossViT3D | sMRI | 0.741 | 0.743 | 0.71 | 5.1M |
| MLP | sFNC | 0.742 | 0.741 | 0.73 | 0.8M |
| ViT | sFNC | 0.750 | 0.745 | 0.72 | 2.6M |
| Multimodal |  | 0.784 | 0.780 | 0.773 |  |
| 3DCNN-CNN | sMRI+sFNC | 0.791 | 0.784 | 0.771 | 11.3M |
| 3DViT-ViT | sMRI+sFNC | 0.782 | 0.779 | 0.774 | 12.3M |
| 3DViT-MLP | sMRI+sFNC | 0.780 | 0.778 | 0.773 | 10.1M |
| **MultiViT** | **sMRI+sFNC** | **0.831** | **0.833** | **0.840** | **10.9M** |

**Fig. 3.**
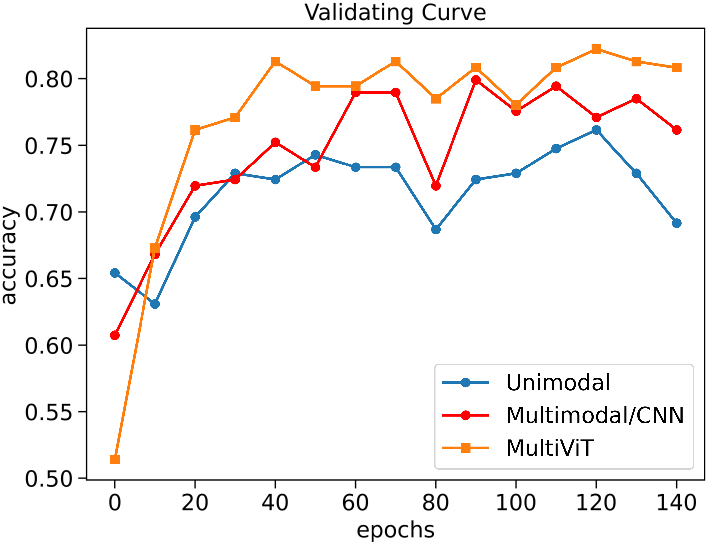
The validation accuracy curves of average(one-model baselines), average(multimodal baselines), and MultiViT.

### C. Discriminative Brain Regions Discovery

#### Attention maps for structural imaging

In this study, we employed the rollout technique [38] to generate attention-based saliency maps, offering an interpretable visualization that identifies specific locations on which the model concentrates. Attention maps can typically discern critical aspects of a model concerning its predictions. We utilized a one-sample t-test to replace the gray matter in each voxel with t-values. We used the testing dataset to ascertain the brain regions contributing most to schizophrenia prediction. Furthermore, we conducted a two-sample t-test between the schizophrenia and healthy control groups to identify vulnerable brain regions associated with schizophrenia. Figure 4 displays the findings of the one-sample t-test for schizophrenia and healthy groups, along with the significant brain regions on which the Multi-ViT model focused to predict outcomes, suggesting potential structural relevance. Some brain regions are more likely to be associated with schizophrenia than healthy controls, while others are less likely to be related to schizophrenia than healthy individuals (Figure 5).

**Fig. 4.**
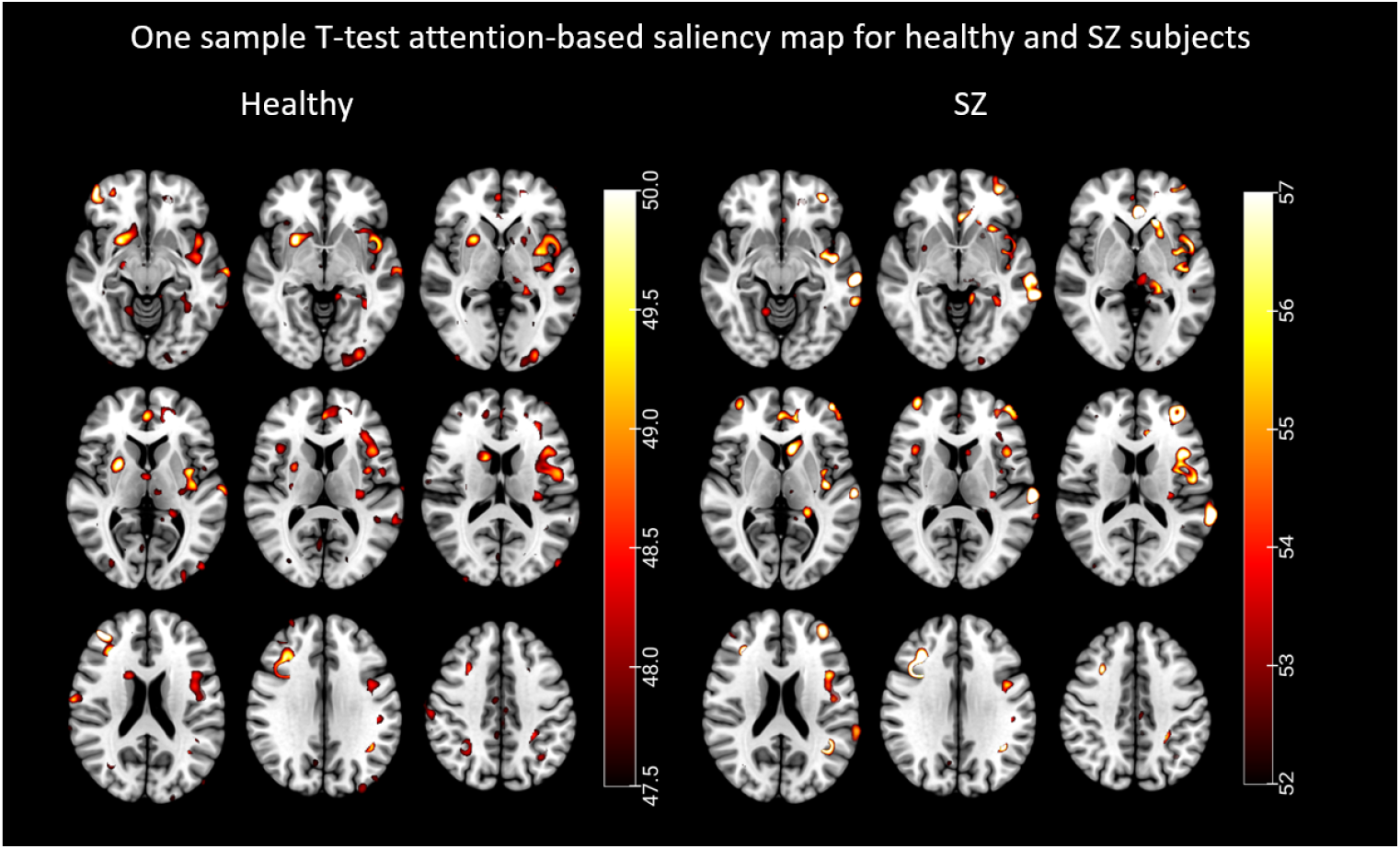
The attention map for one sample t-Test shows highlighted brain regions in which the ViT model focused more. On the left are brain regions more relevant to healthy individuals, and on the right are brain regions more relevant to schizophrenia individuals.

**Fig. 5.**
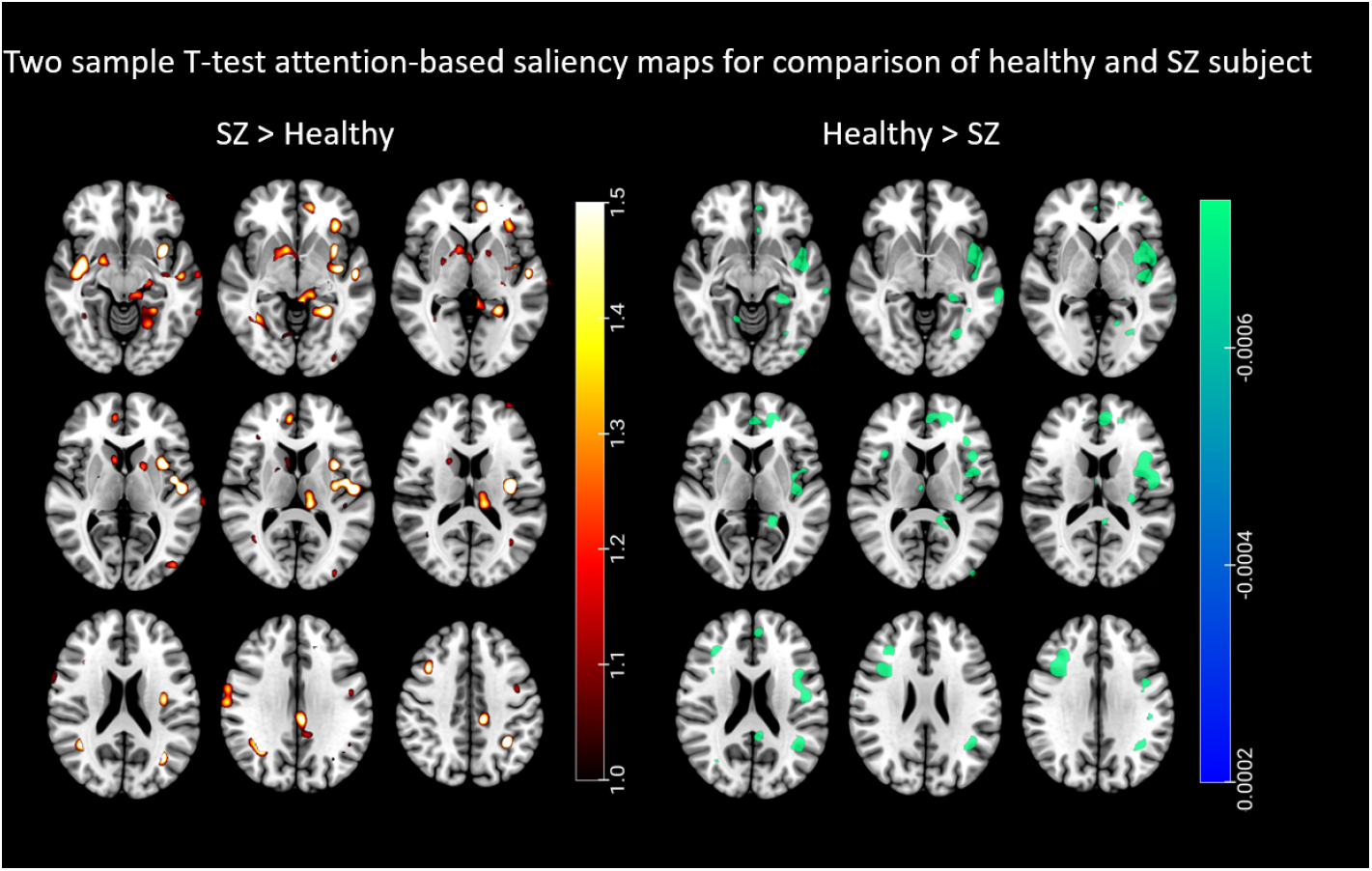
The attention map for the two-sample t-test with a p-value of 0.02 highlights relevance levels of brain regions associated with healthy individuals and schizophrenia. On the left are brain regions in which schizophrenia is significantly more prevalent than in healthy individuals, indicating that these regions are significantly associated with schizophrenia. Right, are the brain regions more relevant to healthy individuals than schizophrenia patients, that is, these regions have a greater significance in the healthy group.

The one-sample t-test results indicate that the schizophrenia and healthy groups share major brain areas, which may overlap. The significant brain regions contributing to the MultiViT model for predicting schizophrenia include the anterior cingulum, lingual gyrus, the middle part of the orbital frontal gyrus, precentral gyrus, insula, cerebellum, supplementary motor area, and hippocampus. The brain regions contributing to healthy predictions encompass the caudate nucleus, superior frontal gyrus, fusiform, lingual gyrus, supplementary motor area, the posterior crus II cerebellum, precuneus, precentral gyrus, and superior temporal gyrus. However, the one-sample t-test cannot distinguish the core regions associated with schizophrenia from those of the healthy group. Consequently, we employed a two-sample t-test to compare the attention maps of individuals from each testing set between the schizophrenia and healthy groups.

The two-sample t-test results on schizophrenia and healthy groups reveal that some brain regions are more significant in the schizophrenia group than the healthy group, including the left cerebellum, left lingual gyrus, middle temporal gyrus, inferior temporal gyrus, and caudate nucleus. Regions more strongly associated with the healthy group include the left precuneus, right cerebellum, left angular gyrus, right inferior frontal gyrus, left caudate nucleus, and right insula.

#### Attention maps for functional connectivity

The FNC matrix represents the temporal correlations amongst spatially distant neurophysiological phenomena. This matrix is commonly employed to evaluate the associations between disparate brain regions and their interactions during diverse cognitive tasks or in a resting state. In the present investigation, we have incorporated the FNC matrix with attention maps derived from the MultiViT model, facilitating the identification of brain functions that the MultiViT model emphasizes during prediction. These functions may strongly associate with schizophrenia and healthy control subjects. Figures 6 and 7 illustrate the attention-based saliency map of the FNC matrix on schizophrenia and healthy control groups. Nevertheless, discerning the distinctions between schizophrenia and healthy control subjects within each group remains challenging. To discern the differences between schizophrenia and healthy subjects, attention-based saliency maps were constructed for the disparities between the schizophrenia and healthy control groups and the inverse. Figure 8 displays the FNC matrix capable of revealing brain functions predominantly associated with schizophrenia patients rather than healthy controls. The labels within the FNC matrix denote various brain functions corresponding to distinct connectivities. The regions exhibiting robust connectivities encompass SC-SM, SC-VS, CB-SM, CB-VS, CC-CC, and VS-DM, which could show potential relevant functions related to schizophrenia.

**Fig. 6.**
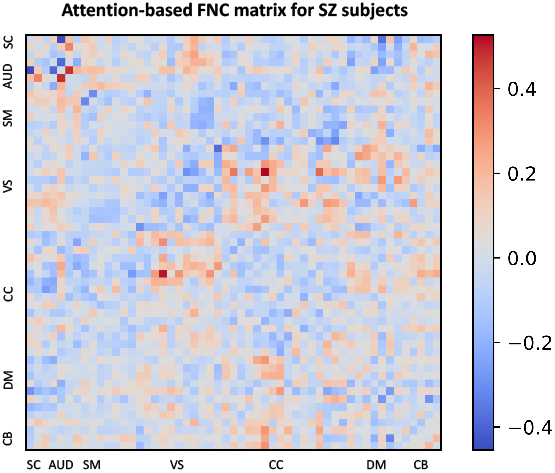
Attention-based FNC matrix for the schizophrenia patients

**Fig. 7.**
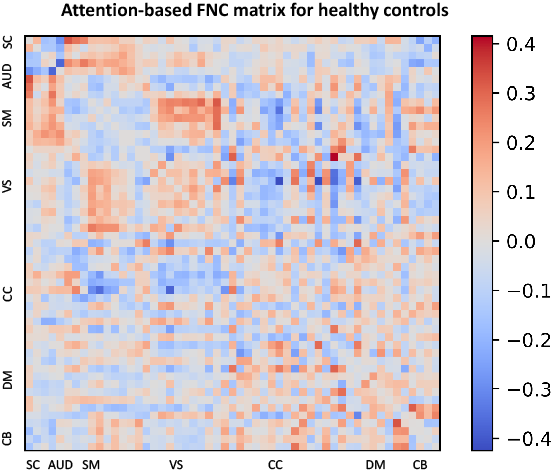
Attention-based FNC matrix for healthy controls.

**Fig. 8.**
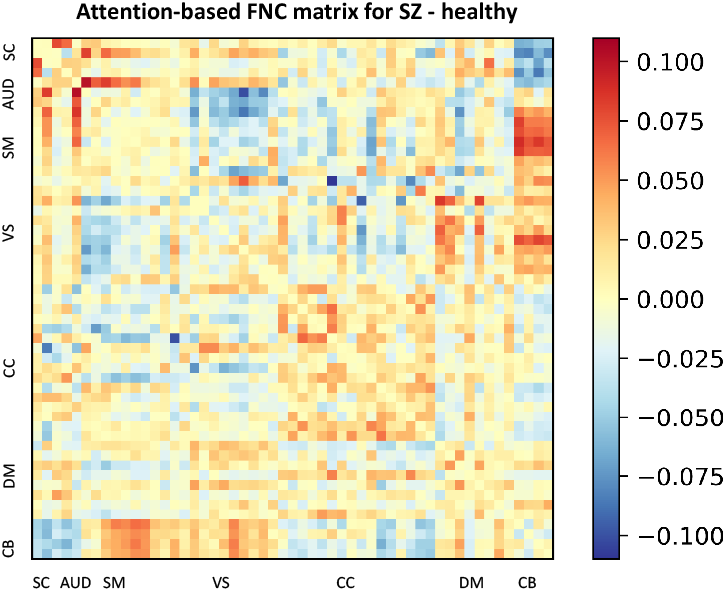
Attention-based FNC matrix for the difference between schizophrenia over healthy controls.

The subcortical (SC) and sensorimotor (SM) interaction plays a pivotal role in subcortical structures, notably the basal ganglia, and thalamus, that are fundamental to the sensori-motor system. The basal ganglia are responsible for initiating and regulating movement by integrating information from the motor cortex and other cortical regions, ultimately promoting seamless and coordinated movement. As a relay center for sensory and motor information, the thalamus facilitates communication with the primary motor and somatosensory cortices to synchronize and modulate motor activity. The Subcortical (SC) and ventral stream (VS) interaction primarily involves the processing of visual information and its integration with other cognitive and motor functions. The pulvinar, a subregion of the thalamus, plays a critical role in this interaction as it relays visual information and communicates with various regions of the visual cortex involved in the ventral stream. Interactions between the cerebellum (CB) and the sensorimotor (SM) network are vital for motor control, coordination, learning, and some cognitive aspects. This interaction ensures precise and well-coordinated movements and the ability to learn and refine new motor skills. Furthermore, the interaction between the cerebellum (CB) and the ventral stream (VS) demonstrates visual-motor integration, wherein the cerebellum can refine and adjust ongoing movements based on visual feedback. Additionally, the cerebellum is involved in visual perception and cognition aspects, such as processing visual motion, estimating time intervals, and predicting future events based on experience. The interaction between different regions of the cingulo-opercular cortex (CC) can involve integrating various cognitive functions, such as attention, working memory, and error detection. Finally, the interaction between the ventral stream (VS) and the dorsal medial prefrontal cortex (DM) involves integrating visual and self-referential information to support various cognitive processes, such as social cognition, perspective-taking, and theory of mind.

The subcortical structures (SC) and cerebellum (CB) interaction plays an essential role in integrating sensory, motor, and cognitive information, which supports motor control, balance, and cognitive processing. This interaction enables the accurate and coordinated execution of movements, balance and posture maintenance, and sensory and cognitive information processing. In the context of schizophrenia, studies have demonstrated impaired connectivity within the brain, especially within connector hubs, including those in the cerebellum and subcortical regions. This impairment has been found to affect subcortical and cerebellar regions and the regions involved in visual and sensorimotor processing [39]. The interaction between the ventral stream (VS) and the sensorimotor (SM) network is crucial for integrating visual and motor information to support perception, action, and cognition. This interaction enables the accurate and coordinated execution of movements, the integration of sensory and visual feedback, and the control of attention and working memory. In the case of schizophrenia, functional disintegration between sensory and cognitive processes has been observed. This disintegration is evident in alterations of both amplitude and connectivity within sensory networks, including within-sensorimotor and sensorimotor-thalamic connections. Sensory nodes also display widespread alterations in the connectivity with higher-order nodes [40]. Finally, the interaction between the ventral stream (VS) and the cingulo-opercular cortex (CC) integrates visual and cognitive information to support attention, decision-making, and cognitive control. This interaction enables the efficient and effective allocation of attention, the integration of reward-related information with decision-making processes, and the flexible adjustment of cognitive processes in response to changing task demands. The connectivity within these regions is also affected in individuals with schizophrenia, characterized by hypoconnectivity between cingulo-opercular regions and hyperconnectivity between the thalamus and sensory cortices. These altered connectivity patterns in schizophrenia highlight the need for a comprehensive, data-driven approach to understanding the complex neuropathology of this disorder [41].

## V. Discussion and Conclusion

In this study, we introduced and applied an innovative multimodal vision transformer pipeline, which not only generates reliable predictions for schizophrenia using clinical datasets but also identifies relationships between two modalities. We successfully experimented with sFNC data as a computational resource for a deep learning model that attains comparable performance to fMRI data after rigorous training [42][43]. Results also indicate that structural and functional changes in schizophrenia have been previously investigated. Karlsgodt et al. [44] analyzed structural MRI and diffusion tensor imaging to demonstrate that schizophrenia patients exhibit reduced gray matter volumes in the medial temporal, superior temporal, and prefrontal areas, which aligns with our findings. Concurrently, they uncovered functional alterations in schizophrenia, revealing that short-term memory and decision-making are more impaired than normal. DeLisi et al. [45] have consolidated multiple findings in schizophrenia, including the cerebellum and superior temporal gyrus.

In our exploration utilizing the interpretable MultiViT model, we pinpointed not only salient structural brain regions associated with schizophrenia but also elucidated potential functional connectivities between various brain regions and their critical functions. For instance, our research unveiled the left cerebellum as a significant brain structure in schizophrenia patients, implicated in movement-related brain functions. Moreover, our FNC analysis showed that the cerebellum exhibits numerous potential functional relationships of considerable importance in schizophrenia patients, encompassing visual-motor integration, motor control, and motor cognition. Additionally, we identified vital brain regions, such as the lingual gyrus, associated with visual and word processing. We demonstrated they possess multiple potential functional relationships with working memory, attention, and cognitive processes. Furthermore, the saliency maps in our study reveal a heightened significance of the middle temporal gyrus among patients with schizophrenia. This brain region plays an essential role in visual and linguistic comprehension and social cognition processes. These findings are congruent with the results from the FNC investigation, which demonstrated a notable interaction correlation between the ventral stream and the dorsal medial prefrontal cortex.

The proposed model is versatile and can be expanded to incorporate an arbitrary number of modalities, theoretically and practically. Our model is expandable and can accommodate additional modalities, as the cross-attention method can link more than two inputs and exchange information effectively. The ViT model, which can handle high-dimensional images and time-series data, is very convenient for training multimodal datasets and through this may have a role to play in helping us understand complex mental illness through the lens of multimodal data fusion.

## Notes

### Competing Interest Statement

The authors have declared no competing interest.

